# Estimating DNA-DNA interaction frequency from Hi-C data at restriction-fragment resolution

**DOI:** 10.1101/377523

**Authors:** Christopher JF Cameron, Josée Dostie, Mathieu Blanchette

## Abstract

Hi-C is a popular technique to map three-dimensional chromosome conformation. In principle, Hi-C’s resolution is only limited by the size of restriction fragments. However, insufficient sequencing depth forces researchers to artificially reduce the resolution of Hi-C matrices at a loss of biological interpretability. We present the Hi-C Interaction Frequency Inference (HIFI) algorithms that accurately estimate restriction-fragment resolution Hi-C matrices by exploiting dependencies between neighboring fragments. Cross-validation experiments and comparisons to 5C data and known regulatory interactions demonstrate HIFI’s superiority to existing approaches. In addition, HIFI’s restriction-fragment resolution reveals a new role for active regulatory regions in structuring topologically associating domains.

Availability: https://github.com/BlanchetteLab/HIFI

## Background

Cells are complex, dynamic environments that require constant regulation of their genes to ensure survival. The advent of chromosome conformation capture (3C) technologies (1), and recent advances in imaging techniques (2), have led to an improved understanding of genome organization and its role in gene regulation (3, 4). Hi-C (5), a high-throughput derivative of 3C, provides an unparalleled view of three-dimensional (3D) genome organization by capturing all DNA-DNA contacts found within a population of cells. Hi-C has revealed different levels of genome organization, including the topologically associating domains (TADs (6, 7), subTADs (8, 9)), and chromatin compartments (5). Yet, the potential for a more refined understanding of 3D genome organization remains largely untapped (10).

In a Hi-C experiment, cross-linked chromatin is digested into fragments using a restriction enzyme (RE). Restriction fragments (RF) are then proximity-ligated to obtain a library of chimeric circular DNA. Paired-end sequencing and mapping of reads to a reference genome identifies interacting RFs and their frequency count. The data is conventionally stored as a pairwise read count matrix, *RC*, where *RC_i,j_* is the number of observed interactions (read-pair count) between genomic regions *i* and *j*. Despite the great sequencing depth of typical Hi-C experiments (200-500 million read pairs), RF-resolution *RC* matrices are extremely sparse, with most RF pairs being observed either zero or one time. This sparsity makes measurements of individual interaction frequencies (IF) between RF pairs inherently stochastic and unreliable. Increasing sequencing coverage is a partial solution, but without improved bioinformatics analyses, the depth of sequencing needed to make reliable estimates of IFs for individual RFs is unmanageable. For this reason, Hi-C data is rarely studied at RF resolution, but instead binned at fixed intervals (e.g., every 25 kb). Unfortunately, reducing the resolution of a Hi-C IF matrix leads to difficulties in studying interactions between fine-scale genomic elements such as promoters and enhancers.

To improve the resolution of Hi-C data, recent protocols suggest digesting DNA more finely, either with a four-cutter RE (10, 11) or DNAse I (12), followed by binning at 1 to 5 kb. While these methodologies increase the resolution of a Hi-C IF matrix, they actually worsen the problem of sparsity and stochastic noise. For example, using a 4-cutter RE instead of a 6-cutter results in a 16-fold increase in the number of RFs, and a 256-fold increase in RF pairs. This problem can be alleviated by using DNA capture technologies to concentrate sequencing on a predefined set of loci (13, 14), but this approach loses the ability to interrogate the whole-genome conformation in a hypothesis-free manner. Instead, new bioinformatics approaches have been proposed to detect individual significant contacts at high resolution from Hi-C data (15, 16), and a machine-learning method has been introduced to smooth Hi-C matrices at 10 kb resolution (17). Dynamic binning was also proposed as a way to adjust bin size to ensure even read coverage across the genome, enabling locally higher resolution (18). However, no approach currently exists to obtain complete and accurate IF matrices at RF resolution. Such an approach would be valuable as it would allow researchers to revisit existing datasets and get more information out of them without having to change experimental protocols or generate more experimental data.

Here, we introduce the Hi-C Interaction Frequency Inference (HIFI) algorithms, a family of computational approaches that provide reliable estimates of IFs at RF resolution. HIFI algorithms reduce stochastic noise, while retaining the highest-possible resolution, by taking advantage of dependencies between neighboring RFs. We validate these algorithms via cross-validation and a comparison to observations made by independent chromosome conformation assays. We further demonstrate that HIFI improves the detection of contacts between promoters and enhancers. Finally, we illustrate additional benefits of high-resolution Hi-C data analysis by using it to study how active regulatory regions are involved in structuring TADs and subTADs.

## Results

HIFI algorithms aim to reliably estimate Hi-C contact frequencies between all intra-chromosomal pairs of restriction fragments. The output of a HIFI algorithm is an IF matrix per chromosome, where each entry (*i,j*) corresponds to the IF of RFs *i* and *j*. As REs do not digest DNA uniformly along the genome, different rows/columns correspond to regions of different sizes. Depending on the RE used, the achievable resolution of Hi-C ranges on average from 434 bp (for a four-cutter such as MboI) to 3.7 kb (for a six-cutter such as HindIII). The high-resolution analysis of Hi-C data faces multiple challenges, of which the sparsity of the observed read-pair data is the most significant. For example, a Hi-C experiment with a very high sequencing depth of one billion read pairs will yield on average approximately 0.1 read pairs per intrachromosomal matrix entry for a 6-cutter RE, and less than 0.001 for a 4-cutter RE. This sparsity results in the observed read-pair count for a given RF pair being a poor (high-variance) estimator of the true IF, except for rare RF pairs located in regions of the Hi-C contact map where IF values are extremely high. All existing solutions to this problem, including the methods introduced in this paper, take advantage of the fact that IFs of neighboring entries in the IF matrix are strongly correlated. In particular, the most common approach to the resolution/accuracy trade off is to artificially reduce the resolution by binning the raw data to fixed-size intervals (e.g., 25 kb bins). This lower resolution increases the number of reads per bin pair, and thus allows for a more reliable estimation of IF, but at the cost of a loss in biological interpretability. Importantly, no unique bin size is uniformly ideal for an entire IF matrix. Portions of an IF matrix where high IFs are present could support a high-resolution analysis, whereas others, corresponding to lower IF values, may require larger bins for accurate IF estimation.

More specifically, the problem addressed here is the following: consider a Hi-C dataset *H* produced with a given restriction enzyme *e*. For a given chromosome, the raw outcome is stored in an *n × n* intrachromosomal matrix *RC*, where *n* is the number of RFs produced by *e*, and *RC_i,j_* contains the number of read pairs mapped to RF pair (*i,j*). Our goal is to estimate as accurately as possible the true RF-level interaction frequency matrix, *IF_true_,* which is the theoretical *n × n* IF matrix one would obtain if one were to sequence an infinitely large version of *H* to infinite depth (scaled for the total number of read pairs). *IF_true_* is affected by a number of library, sequencing, and mapping biases that would need to be corrected in order to allow for proper biological interpretation; many such normalization techniques already exist for this task (19–21). Our goal here is not to improve upon these techniques, but to work upstream and provide the most accurate estimate of *IF_true_*.

Four approaches are introduced and assessed (see Methods for details), each taking as input matrix *RC* and producing as output an estimate of *IF_true_*:

1. The commonly-used fixed-binning approach, where the genome is first partitioned into bins containing a fixed number of kb (or a fixed number of RFs) and the estimated IF for a given RF pair is the average of the *RC* values of all RF pairs that belong to the same bin pair.
2. A simple Kernel Density Estimation (HIFI-KDE) approach, where the IF estimate at a given matrix entry is obtained as the average of surrounding entries, weighted using a two-dimensional Gaussian distribution with a fixed standard deviation (bandwidth).
3. An Adaptive Kernel Density Estimation (HIFI-AKDE) approach, where the bandwidth is chosen dynamically for each matrix entry in order to ensure that a sufficient number of read pairs is available for reliable IF estimation, while maximizing the resolution.
4. An approach based on Markov random fields (HIFI-MRF) where dependencies between neighboring cells are modeled and used to identify the maximum *a posteriori* estimate of *IF_true_*.

Assessing the accuracy of high-resolution IF inference algorithms is challenging because *IF_true_* is unknown, as Hi-C datasets of infinite sequencing depths are not achievable. Instead we consider two surrogates. First, we use a crossvalidation approach from existing Hi-C data. Second, we assess predictions against data produced by Chromosome Conformation Capture Carbon Copy (5C (22)), a targeted amplification protocol that achieves a much higher read count per RF pair compared to Hi-C.

### Cross-validation of HIFI algorithms

We used crossvalidation to assess the accuracy of HIFI algorithms genome-wide. Here, a Hi-C read-pair dataset of high sequencing depth produced by Rao et al. (2014) (23) from GM12878 cells using HindIII was first filtered to retain only high-confidence intra-chromosomal read pairs. These read pairs were then randomly partitioned into an input set (containing 80% of the set of filtered read pairs, or 607,587,043 read pairs), and a test set (20%, or 151,979,454 read pairs) (Fig. 1A). The input set is then further downsampled into seven subsets ranging in size from 1 to 100% of the full input set. Mapping and tabulating read pairs at RF-level resolution yields a family of read count matrices: *RC*_*input*_1_, *RC*_*input*_2_,…, *RC*_*input*_100_, and *RC_test_*.

**Fig. 1.**
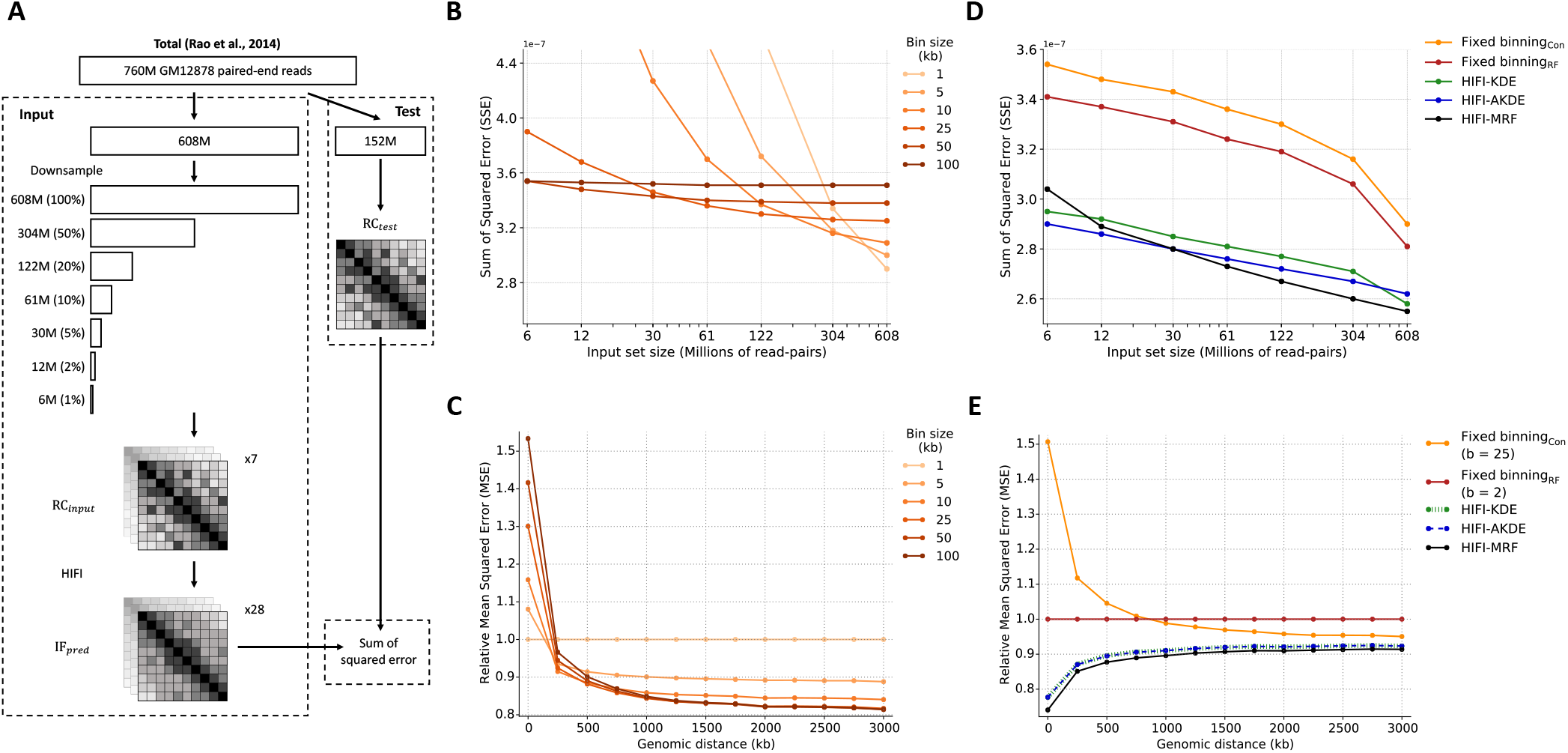
Cross-validation of fixed-binning and HIFI methodologies. **A**) Schematic representation of cross-validation methodology to assess the accuracy of fixed-binning and proposed HIFI methodologies. **B**) Cross-validation error for canonical fixed-binning approaches, for different bin sizes, as a function of coverage. See also Suppl. Fig. 1 for similar analyses for RF fixed binning, HIFI-KDE and HIFI-AKDE. **C**) Analysis of canonical fixed-binning error (relative to error with one RF per bin) across genomic distance between RF-pairs. No singular bin size performs best for all genomic distances. **D**) Comparison of errors for different approaches. For fixed binning and HIFI-KDE, the optimal bin size or bandwidth was chosen separately for each coverage level. Nonetheless, HIFI-MRF outperforms all other approaches. **E**) Comparison of errors (relative to error obtained with fixed binning using 2 RF per bin) by genomic distance of RF pairs, using as input a set of 304M read pairs (50% of total training set). HIFI-MRF performs best across all distances.

Each of the four inference algorithms are evaluated by their application to each of the downsampled input matrices to obtain a predicted IF matrix, *IF_pred_*, which is then compared to the test matrix *RC_test_* to obtain the sum of squared errors:

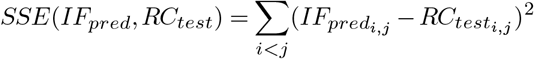

Although *RC_test_* is not equal to *IF_true_*, the inference approach that minimizes *SSE*(*IF_pred_, RC_test_*) is also the one that minimizes *SSE*(*IF_pred_, IF_true_*), and hence, this serves as a valid basis for comparison.

Fig. 1B and Suppl. Fig. 1A show that the accuracy of fixed-binning strategies improves with input set size and that the optimal accuracy is obtained at different bin sizes for different input set sizes: large bins are ideal for low-coverage training data, whereas smaller bins are better with high-coverage data. More importantly, the fact that read pairs are highly non-uniformly distributed in *RC* matrices means that the ideal bin size differs depending on the local *RC* density. In particular, short-range contacts, which typically have higher *RC* values, can support high-resolution analyses (smaller bins), but those at longer ranges are best estimated with larger bins (Fig. 1C, and also Suppl. Fig. 1B for similar analyses for binning based on number of RFs rather than sequence size). The HIFI-KDE approach with a fixed bandwidth generally obtains better results (Fig. 1D and Suppl. Fig. 1C), but suffers from the same type of problem, where optimal results are obtained with large bandwidth values for low-coverage regions and lower bandwidth values for high coverage regions. The HIFI-AKDE approach, where different bandwidth values are chosen at each cell based on the surrounding signal density, outperform the first two approaches (Fig. 1D and Suppl. Fig. 1D), with optimal performance obtained using a *MinimumCount* value of 100 (see Methods) throughout various coverage levels. HIFI-MRF performs the best overall (Fig. 1D and Suppl. Fig. 1E), except at extremely low sequencing depths (i.e., 6-12M read pairs). Indeed, for typical sequencing depths (100-250M read pairs), HIFI-MRF improves IF estimation accuracy over the entire range of genomic distances (Fig. 1E) producing estimates that are 5-40% more accurate than those obtained by fixed-binning approaches and 5% more accurate than HIFI-KDE and HIFI-AKDE. See Suppl. Fig. 2 for an example of HIFI-MRF-processed HiC matrix and a comparison to fixed-binning analysis. We also attempted to use the same strategy to evaluate HiPlus (17), a machine-learning technique for high-resolution analysis of Hi-C data, but found that the model did not perform well on non-bias corrected Hi-C data for this analysis.

**Fig. 2.**
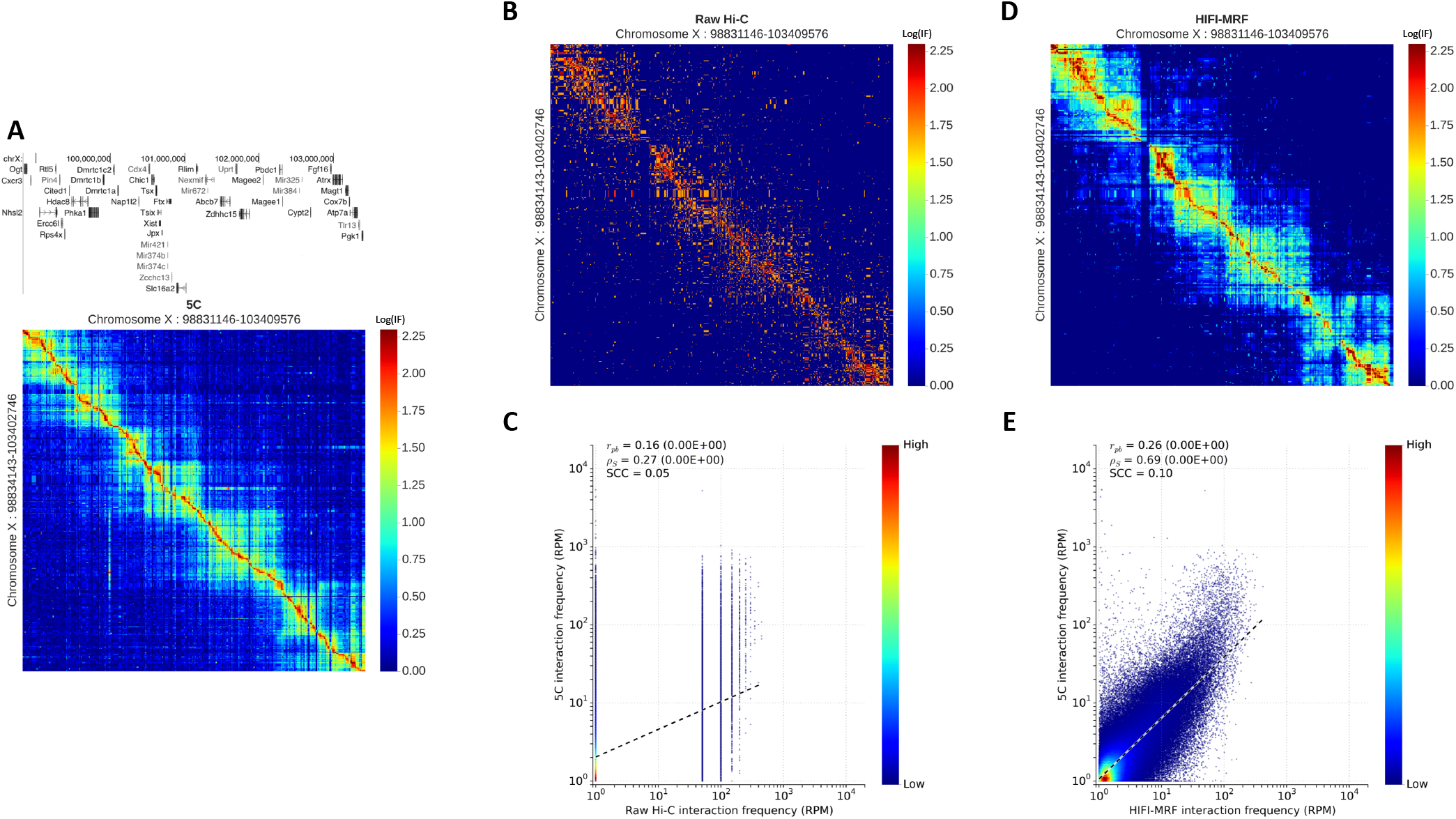
Recapitulation of 5C observations by HIFI-MRF. **A**) IF matrix obtained by 5C of the 4.5 Mb locus surrounding the *X_ist_* gene in mouse embryonic stem cells (7). Note the use of true-size heatmaps, where the height (resp. width) of a row (resp. column) is proportional to the size of the RF it represents. **B**) Raw, RF-resolution Hi-C data for the same region (6). **C**) Correlation of 5C and raw Hi-C data at RF resolution (Pearson *r_pb_* = 0.16, p-value< 10^−16^; Spearman *ρ_s_* = 0.27, two-sided Student’s *t*-test p-value< 10^−16^); stratum-adjusted correlation coefficient (SCC) (29) = 0.05). **D**) IF matrix estimated by HIFI-MRF from the same Hi-C data. Observe the similarity to the 5C data in (**A**), **E**) Correlation of 5C and HIFI-MRF processed HI-C data at RF resolution (Pearson *r_pb_* = 0.26, p-value< 10^−16^; Spearman *ρ_s_* = 0.69, p-value< 10^−16^); SCC = 0.10).

### Validation against 5C data

5C has been used to study the conformation of moderate-size genomic regions (100 kb - 5 Mb), including the beta-globin locus (22, 24), the *HOX* clusters (8, 25, 26), the *CFTR* locus (27, 28) and the *Xist* locus (7). 5C allows for a high sequencing depth measurement of the IF of each RF pair within given genomic regions, which improves the accuracy of RF-level IF estimates. As such, 5C data constitutes an excellent benchmark to compare different inference approaches. We analyzed data from two cell types for which both 5C and Hi-C data are available: (i) a 4 Mb region around the *Xist* gene (Fig. 2A,B) in mouse embryonic stem cells (mESC; Hi-C data from Dixon et al., 2012 (6); 5C data from Nora et al., 2012 (7)), and (ii) a 2.7 Mb region around the *CFTR* gene (Suppl. Fig. 3A,B) in human GM12878 cells (5C data from Smith et al., 2016 (28); Hi-C data from Rao et al., 2014 (23)). In the GM12878 dataset, which has higher Hi-C sequencing depth (760M mapped read pairs genome-wide), the correlation between raw Hi-C and 5C data is moderate (Spearman *ρ_s_* = 0.45; Suppl. Fig. 3C), but it is improved by the application of HIFI-MRF (*ρ_s_* = 0.71; Suppl. Fig. 3D,E and Suppl. Fig. 4). In the mESC dataset, with lower Hi-C sequencing coverage (122M read pairs), the correlation of raw 5C against raw Hi-C data is relatively weak (*ρ_s_* = 0.27; Fig. 2C), but improves to nearly the same level as in the first dataset from the application of HIFI-MRF (*ρ_s_* = 0.69; Fig. 2D,E and Suppl. Fig. 5). Strata-adjusted correlation coefficients (SCC) (29), which factor out correlations induced by genomic distance dependencies, are also improved by the application of HIFI-MRF. Also note how, in both cases, the intricate structure of TADs, as well as some of the finer looping events become apparent in the HIFI-MRF-processed Hi-C data (Fig. 2D and Suppl. Fig. 3D).

Indeed, the application of HIFI-MRF to Hi-C data allows for the detection of regulatory contacts that could previously only be observed using 5C. For example, Nora et al. (2012) (7) used 5C to observe a long-range interaction between *Tsix* and its transcriptional regulator – a large intervening non-coding RNA called “Linx” – occurring in female mice as a component of X-inactivation. This interaction is very clearly observed in the HIFI-MRF-processed Hi-C data (Dixon et al., 2012) (6), whereas it is difficult to distinguish from background in raw or binned Hi-C data (Fig. 3A, 3B). These results demonstrate that HIFI-MRF can be used to analyze existing Hi-C data sets and potentially lead to novel discoveries at finer genomic scales.

**Fig. 3.**
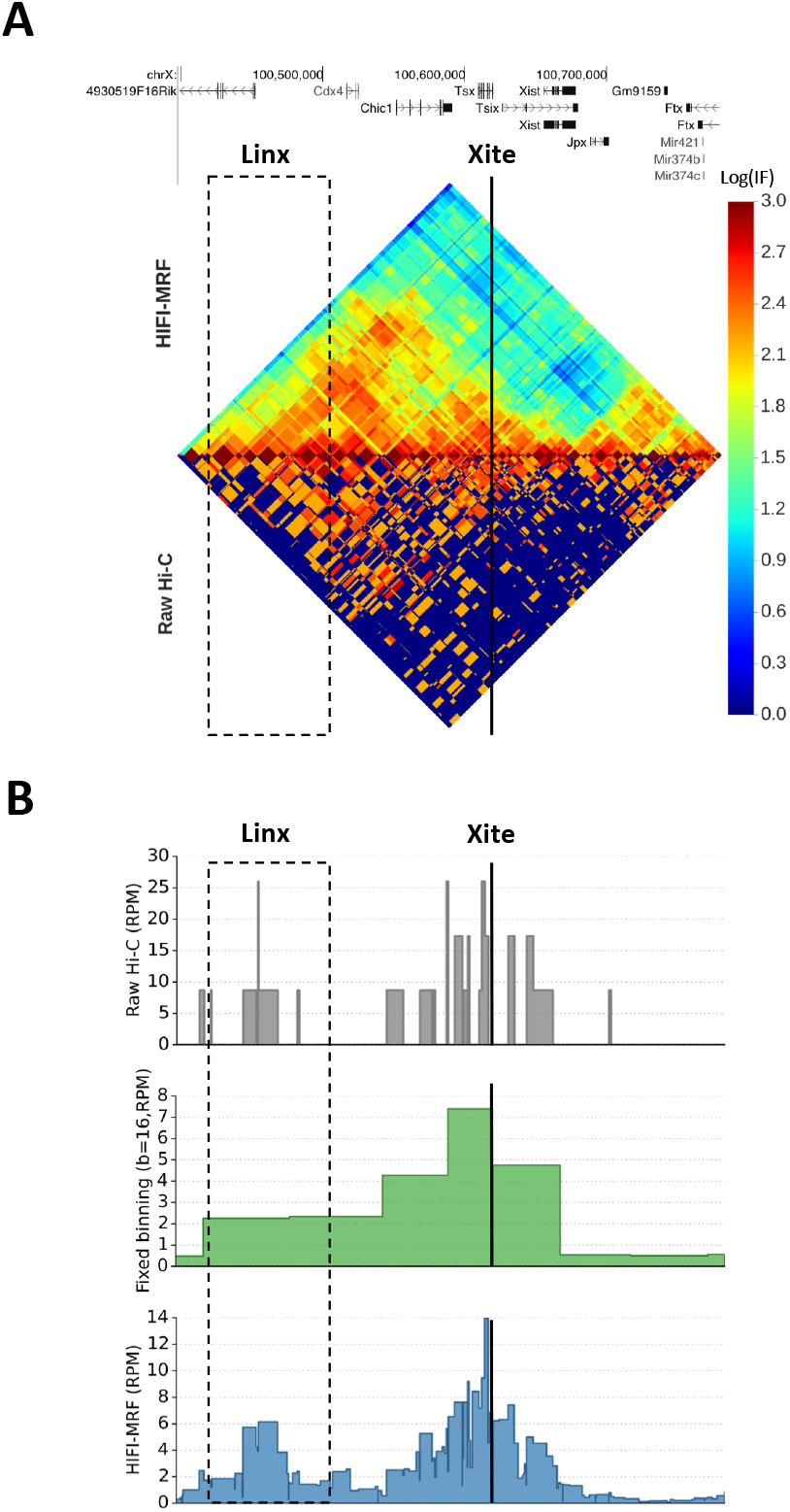
HIFI-MRF reveals fine-scale regulatory contacts in Hi-C data. Heatmap (**A**) and virtual 4C (53, 54) plots based on raw, binned, or HIFI-MRF processed data (**B**) showing the long-range interaction between *Tsix* and its transcriptional regulator, *Linx*, on chromosome X of female mice as observed by Nora et al. (2012) (7) using 5C. This interaction is more easily observed in HIFI-MRF data than in raw or binned Hi-C data.

### Validation against externally-predicted chromatin contacts

To more fully assess the extent to which HIFI-MRF-processed Hi-C data can be used to identify biologically relevant contacts, we asked whether it can also confirm chromatin interactions found through alternative approaches. Specifically, we considered a set of contacts identified by Chromatin Interaction Analysis with Paired-End Tag Sequencing (ChIA-PET (30)) in GM12878 cells, bound either by CTCF (92,114 contacts (31)), RNA Polymerase II (PolII - 192,394 contacts (31)), or RAD21 (38,952 contacts (32)). We also considered a set of computationally inferred contacts identified by correlation of DNAse I hypersensitivity signals across multiple cell types (33). For each set of contacts, a set of negative (control) fragment pairs were chosen by randomly re-pairing the same RFs. We then measured, for each range of genomic distance, the extent to which positive contacts could be distinguished from negative contacts on the basis of normalized HIFI-MRF Hi-C data, by measuring the Area Under the Receiver Operating Characteristic curve (AUROC) of a univariate predictor using the RF pair’s inferred IF value as a predictive variable. Higher AUROC values indicate improved ability to distinguish positive from negative contacts. We observe that HIFI-MRF-processed Hi-C data allows significantly better detection of validated contacts compared to fixed-binning approaches, for all four datasets, across all genomic distance ranges, and both at low (61M read pairs obtained by downsampling; Fig. 4A-C and Suppl. Fig. 6A-C, 7A,B), and high (608M read pairs; Fig. 4D-F, Suppl. Fig. 6D-F, 7C,D) sequencing depths. Notably, the ability to distinguish positive from negative ChIA-PET contacts is relatively poor at short distances (<50 kb) because nearly all pairs have very high IF values, but improves considerably at longer range (300-500 kb). In contrast, contacts inferred based on DHS correlations are more difficult to identify overall (AU-ROC<0.6), becoming increasingly so at longer ranges. We speculate that this loss in detection power may be due to an increased error rate present in this benchmark dataset. Remarkably, the application of HIFI-MRF to low-coverage Hi-C data yields predictive power that is nearly as good as in the high-coverage dataset (compare panels Fig. 4A-C to 4D-F), suggesting that HIFI-MRF is able to identify functional contacts even in Hi-C data of moderate depth. Fig. 4 also includes results for HiCPlus (17) and HMRFBayes (15), an approach for the detection of significant contacts at RF resolution (see Methods). Overall, HIFI-MRF clearly outperforms these two approaches, although HMRFBayes performs nearly equally well for some low-coverage data sets (Fig. 4A-C). The advantage of HIFI-MRF is particularly noticeable at short-to medium-range distances (<200 kb). Taken together, these results show that using HIFI-MRF to process Hi-C data improves the ability to delineate individual chromatin contacts.

**Fig. 4.**
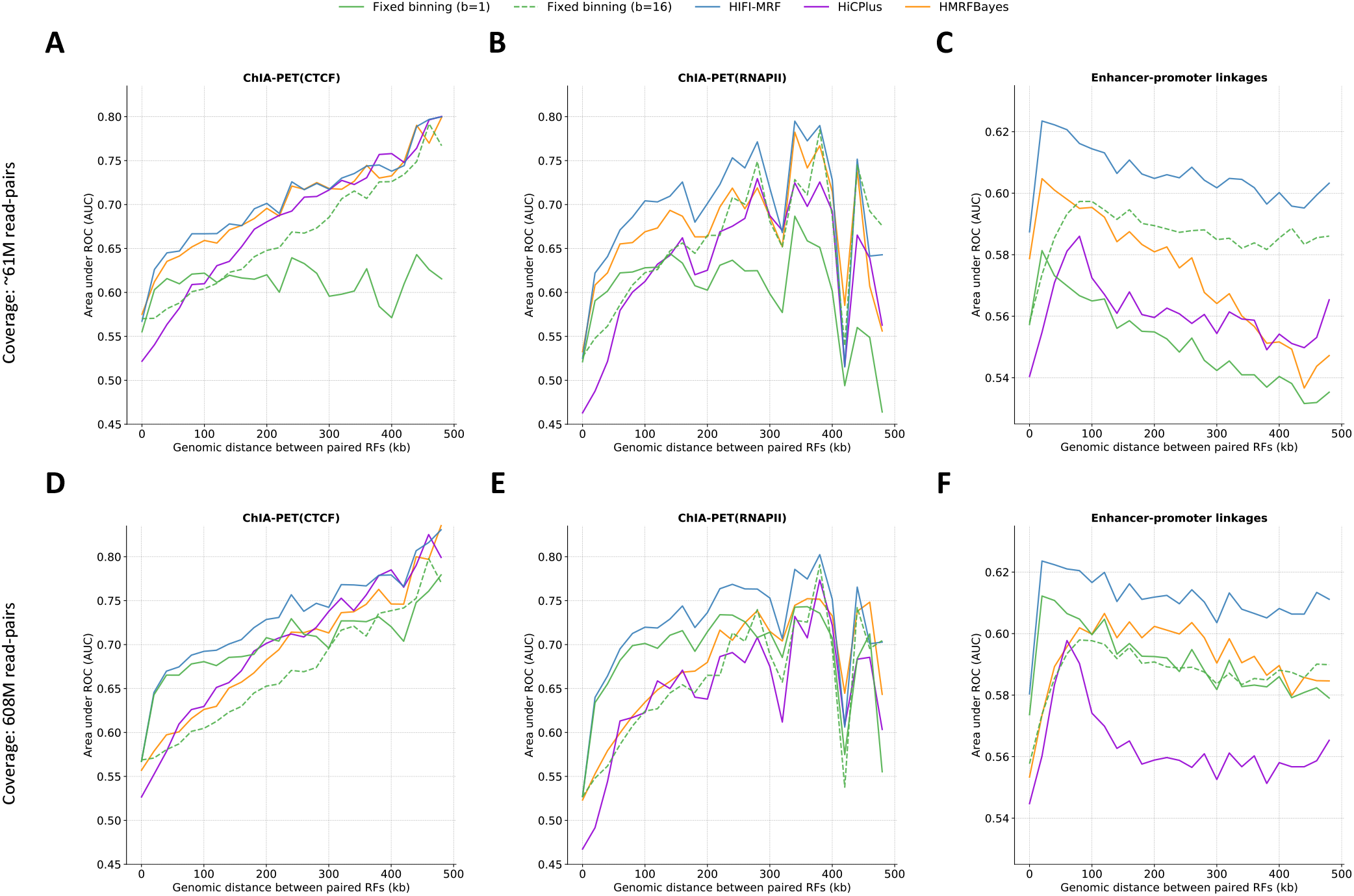
Positive/negative RF contact delineation analysis. The ability of different HiC data analysis approaches to distinguish positive fron negative (control) contacts is measured, for various data sets, using the area under the receiver operating characteristic curve (AUROC) for univariate predictors using as input the predicted *IF* values. (A and D) CTCF-mediated contacts identified by ChIA-PET (31); (B and E) RNAPII-mediated contacts identified by ChIA-PET (31); (C and F) Inferred enhancer-promoter linkages based on DHS correlation (33). To allow for the comparison with HiCPlus and HMRFBayes, only contacts occurring on chromosomes 9 to 22, X, and Y, and within a distance of 1 Mb, are analyzed. Top (**A-C**) and bottom (**D-F**) rows represent the performance of the classifiers applied to Hi-C data of size 60.8M (10% of input set) and 608M (100% of input set), respectively. Genome-wide results for HIFI are shown in Suppl. Fig. 6 Similar results are observed for ChIA-PET RAD21 (Suppl. Fig. 7).

### HIFI allows new insight into fine-level genome organization

The high accuracy and resolution afforded by HIFI enables researchers to answer questions that are difficult to address with lower-resolution analyses of Hi-C data. Here, we illustrate one such application: the high-resolution analysis of TAD and subTAD boundaries. We used a modified directionality index (DI) score, originally introduced by Dixon et al. (2012 (6); see Methods), to identify 5,000 TAD boundaries in the HindIII-GM12878 Hi-C data. Boundary predictions were performed at two resolutions: (i) RF-resolution using HIFI-MRF-processed data (3.7 kb on average; Fig. 5A, top heatmap) and (ii) classical fixed-binning approach (16 RF ≈ 50 kb per bin; Fig. 5A, bottom heatmap). Using ENCODE ChIP-seq datasets (34), we quantified the occupancy of DNA-binding proteins relative to TAD boundaries. Consistent with previously reported observations and models (23, 35, 36), CTCF (Fig. 5B) showed a remarkable enrichment immediately outside of these boundaries, with sites on the plus strand sharply peaking at upstream TAD boundaries and those on the minus strand peaking at downstream boundaries. Similar enrichments at TAD boundaries are observed for RAD21, SMC3 (Cohesin complex), YY1, and ZNF143 (Suppl. Fig. 8), consistent with previous reports (6, 37–42). Although the same phenomenon is visible in fixed-binning data, the peaks are much sharper (narrower and higher) in HIFI-MRF data, indicating that RF-resolution allows more accurate calls of TAD boundaries.

**Fig. 5.**
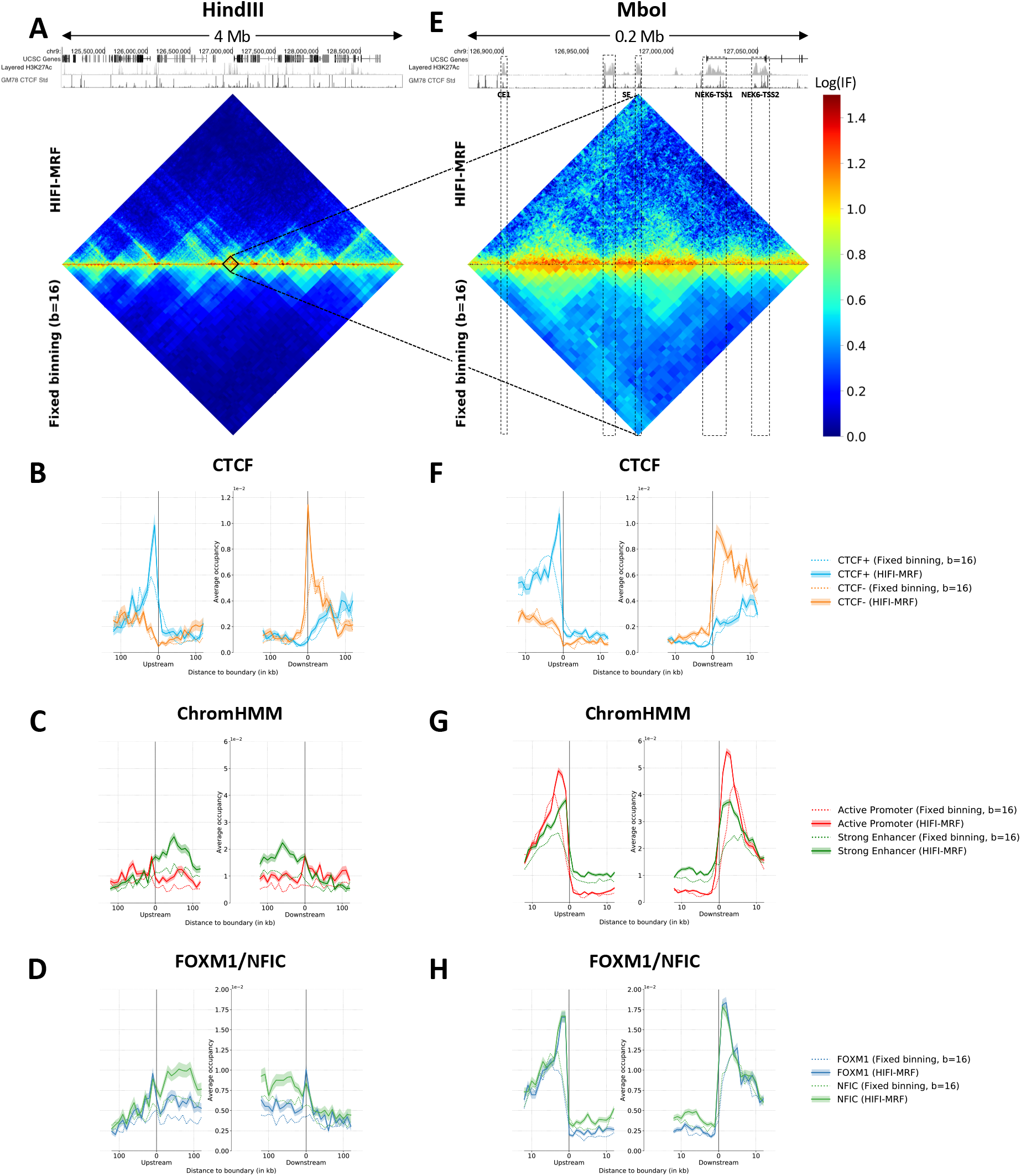
Analysis of RF-resolution TAD and subTAD boundaries in GM12878. Analyses were performed on both HI-C data resulting from a HindIII (3.4 kb per RF on average; panels **A-D**) and a MboI restriction digest (434 bp per RF on average; panels **E-H**), from Rao et al. (2014) (23). TAD and subTAD boundary predictions were made on IF matrices produced either by HIFI-MRF or a fixed-binning approach (16 RF per bin, i.e. approx. 50 kb per bin for HindIII and 7 kb per bin for MboI). **A**) IF matrices produced by HIFI-MRF (top) and fixed binning (bottom) for a 4 Mb locus surrounding the NEK6 locus (chr9:124999244-128993971). (**B** and **F**) CTCF occupancy as a function of distance to the nearest TAD (**B**) or subTAD (**F**) boundary, separately for sites on the forward and reverse strands. Convergent CTCF sites are enriched at both TAD and subTAD boundaries. Shaded band indicate 95% confidence intervals of the estimate of the mean occupancy. **C** and **G**) Coverage of active promoters (red) and strong enhancers (green) identified by ChromHMM, as a function of the distance to the nearest TAD (**C**) or subTAD (**G**) boundary. These regions are very strongly enriched just outside of subTAD boundaries, but less so around TAD boundaries. (**D** and **H**) Occupancy of two transcription factors, FOXM1 and NFIC, as a function of distance to the nearest TAD (**B**) or subTAD (**F**) boundary. While most TFs have an occupancy peak at TAD and subTAD boundaries, the extent of the enrichment within TADs varies from low (e.g. FOXM1) to high (e.g. NFIC). (**E**) IF matrices produced by HIFI-MRF (top) and fixed binning (bottom) for the 200 kb NEK6 locus (chr9:126879748-127079891). Regulatory regions identified in Huang et al. (2016) (55) are marked SE (super enhancer), CE1 (conventional enhancer), NEK6-TSS1 and NEK6-TSS2 (alternative promoters). Notice how all these regions lie between visible subTADs.

We next studied the role of TAD boundaries in gene regulation, by looking at the distribution of active regulatory regions, as annotated by ChromHMM (43) based on cell-type-specific histone marks and DNA accessibility data. We observe a moderate enrichment for active promoters immediately outside TAD boundaries (only visible in HIFI-MRF processed data) and for strong enhancers within TADs. This trend is partially reflected in the occupancy profiles of several transcription factors (Fig. 5D and Suppl. Fig. 9). These transcription factors (in particular EBF1, EP300, IKZF1, MEF2A, MEF2C, and NFIC) exhibit a gradual enrichment toward the middle of TADs, together with a small but well-defined, CTCF-like peak just outside TAD boundaries. While others (e.g., FOXM1, IRF4, and RUNX3) show a more prominent peak at TAD boundaries (Suppl. Fig. 10A-C). Other (e.g., GABPA, MYC, and SIX5) demonstrate a depletion of occupancy within TADs (Suppl. Fig. 10D-F). Notice that in many cases, the enrichment at TAD boundaries is only apparent based on HIFI-MRF data and would likely be missed using data binned at 50 kb resolution.

We then repeated the analysis (HIFI-MRF followed by TAD boundary calls) on Hi-C data generated on the same cell line using the 4-cutter MboI restriction enzyme, with cut sites every 434 bp on average. The extremely high resolution of this dataset (Fig. 5E) provides opportunities to study fine structures such as subTADs (8, 9), which are difficult to study at lower resolutions. We used the HIFI-MRF MboI-GM12878 data and the same modified DI approach to identify a set of 25,000 domain boundaries, of which approximately 2,500 matched a HindIII-GM12878 TAD boundary (within 25 kb). The remaining ~22,500 boundaries are not detected in the HindIII data and likely correspond to subTAD boundaries. Repeating the occupancy analysis against subTAD boundaries, the same enrichment for convergent CTCF sites is observed (Fig. 5F), but a very different picture emerges with respect to regulatory regions. Most notably, active promoters, and to a lesser extent strong enhancers, have a clear tendency to occupy regions that lie immediately outside sub-TADs (Fig. 5G; see also example in Fig. 5E). Indeed, the density of active promoters is approximately 30 times higher in the 1 kb region that precedes a subTAD boundary than in the 1 kb region that follows one. A similar enrichment is found in inter-subTAD regions for FOXM1 and NFIC (Fig. 5H), and nearly all transcription factors studied. These results are consistent with a model where active regulatory regions play a key role in partitioning TADs into subTADs.

## Discussion and Conclusions

Hi-C has become a commonly used approach to map 3D chromatin organization genome-wide. Since its introduction in 2009, the method has been updated many times to improve upon accuracy and resolution, or to target specific types of contacts. However, to date, using Hi-C data to accurately and systematically identify fine-scale chromosome contacts remains challenging, mostly because the sequencing depth required to achieve high-resolution contact maps is too great. To overcome the sparsity of contact information and increase the signal-to-noise ratio, Hi-C data is traditionally binned at fixed intervals along chromosomes to produce lower-resolution matrices (10). This lower-resolution representation of Hi-C data limits its application in studies of genomic regulatory networks or mechanisms of disease, which require robust, high-resolution 3D genomics data.

Here, we introduced HIFI, a family of density estimation algorithms that allow for the observation of high-resolution (at the restriction-fragment scale) genomic contacts from Hi-C data of various sequencing depths. Our results show that HIFI algorithms, and in particular those based on Markov Random Fields (HIFI-MRF), provide highly accurate estimates of Hi-C interaction frequency at RF resolution, and outperform classical fixed-binning approaches. We demonstrate that HIFI-MRF recapitulates contact data obtained by 5C and also captures interactions detected by ChIA-PET (Fig. 4) better than HiCPlus and HMRFBayes (15). Unlike the former, HIFI is easy to use and does not require special equipment (GPUs) to run within a reasonable time frame. HIFI also runs more than 100 times faster than the HMRFBayes. The high resolution and accuracy provided by HIFI allows analyses and discoveries that could not be made with lower-resolution Hi-C data. For example, HIFI allows for the identification of TAD boundaries at RF resolution, which provides a unique opportunity to finely delineate the role of different DNA-binding proteins. Benefiting from the RF-resolution achieved with HIFI-MRF, we show that CTCF, RAD21, SMC3, and ZNF143 are enriched just outside both TAD and subTAD boundaries and their sharp depletion within-TADs may be a major contributor to the formation of TAD boundaries (Fig. 5). In addition, we detail a set of transcription factors (based on ENCODE ChIP-seq data) that are found to be enriched at RFs labeled as TAD boundaries (Fig. 5B,C). Finally, we highlight the new observation that active enhancers and promoters appear to provide structure to TADs, whereby DNA located between consecutive active regulatory regions form subTADs. This is obviously just an illustration of insights that can be gained from the analysis of Hi-C at high resolution. Others would include the use of HIFI-processed Hi-C data to further dissect the mechanisms of genome organization, and to prioritize non-coding variants obtained from genome-wide association (GWAS) or expression quantitative trait loci (eQTL) studies, as is starting to be done with capture Hi-C data (44).

While HIFI provides a significant improvement over previous methodologies for handling Hi-C matrix sparsity, there remains several directions for possible improvements. First, HIFI is relatively slow, requiring roughly an hour per chromosome (at HindIII resolution), due to the size of the matrices analyzed and the complexity of MRF-based inference. Improved algorithms, multi-threading, and GPU-based computation are expected to provide significant speed-ups and are under development. These improvements will also allow the calculation of confidence intervals for estimated contacts frequencies, using Markov Chain Monte Carlo sampling. Machine-learning (ML) approaches, such as convolutional neural nets (CNN), offer an alternative to probabilistic approaches like HIFI-MRF. In recent work by Zhang et al. (2018) (17), the authors showed that CNNs can be trained on Hi-C data to increase the resolution from 40 kb to 10 kb. Being model-free, ML approaches have the potential to discover and take advantage of unsuspected dependencies in the data. However, these models have yet to produce RF-resolution data and thus, remain limited in their ability to provide biological support as shown in this manuscript. In addition, being intrinsically complex models, prediction errors may occur in unexpected manners.

In conclusion, the HIFI algorithms and software described in this manuscript allow for accurate, high-resolution analyses of 3D genome organization using Hi-C data. RF-resolution Hi-C data allows for the recapitulation of observations made by 5C, a better separation of positive and control/background contacts, RF-resolution TAD and subTAD boundary calling, and the identification of potential DNA-DNA contacts and TF enrichments that drive changes in chromatin architecture and gene regulation. By operating upstream of many Hi-C data analysis tools (e.g., loop, TAD, and compartment predictors as well as fragment-bias normalization), HIFI can easily be inserted in a number of Hi-C data analysis pipelines, and we believe that the research community will be quick to take advantage of this family of new algorithms.

## Methods

### Hi-C read-pair pre-processing

The publicly available Hi-C User Pipeline (HiCUP (45)) v0.5.3 was used to process raw sequencing reads. HiCUP-mapped reads to the human (hg19) genome are also filtered to remove expected artifacts resulting from the sonication and ligation steps (e.g., circularized reads, reads with dangling ends) of the Hi-C protocol. Mapped reads were further filtered for a Mapping Quality Score (MAQ) greater than 30 (19). BAM/SAM-mapped read files were then converted (by our ‘BAMtoSparseMatrix.py’ script) to a raw read-pair count matrix *RC*, stored using a sparse matrix TSV file format, before use with HIFI.

### HIFI algorithms

The HIFI package is available at https://github.com/BlanchetteLab/HIFI. It consists of a C++ program for IF estimation, together with Python scripts for input data formatting and the true-size IF matrix visualization. This section provides algorithmic details.

#### Fragment-specific bias calculation

Factors such as fragment size, GC content, and mappability affect the observed read count matrix *RC*. For each fragment *i* of chromosome *c*, we estimate this bias as

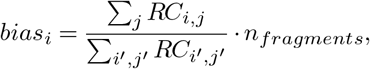

where *n_fragments_* is the number of RFs on that chromosome. Computed biases are used to obtain a normalized read count matrix *nRC*, where 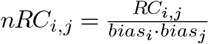.

#### Fixed binning approach

In the fixed-binning approach, the user specifies the value *binSize*, which is the number of consecutive RFs to be binned together. Defining *bin_i_* = {*j*: ⎿*j/binSize*⏌ = ⎿*i/binSize*⏌}, we then obtain the following estimate of interaction frequency for RF pair (*i,j*):

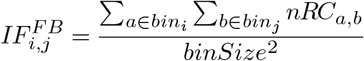

#### Fixed Kernel Density Estimation (HIFI-KDE)

This approach follows the standard two-dimensional Kernel Density Estimation (KDE) procedure (46, 47), where predicted *IF* for RF pair (*i,j*) is obtained as a weighted sum of the entries of *RC* surrounding (*i,j*), parameterized by bandwidth parameter *h*. Specifically, we set

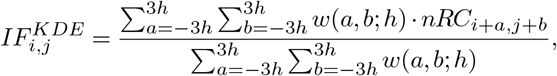

where 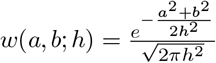. Near the edges of the matrix, values of *a* and *b* such that indices (*i + a, j + b*) fall outside the matrix are excluded from the sums of both the numerator and denominator.

#### Adaptive Kernel Density Estimation (HIFI-AKDE)

This approach is similar to the fixed KDE, except that the value of the bandwidth parameter *h* is chosen separately for each pair (*i,j*). Specifically, we choose *h_i,j_* to be the smallest value such that

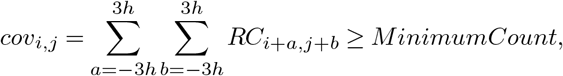

where ‘*MinimumCount*’ is a user-defined parameter. In other words, regions of the matrix that tend to have larger *RC* values are estimated using smaller bandwidths (i.e., higher resolution), whereas those that are more sparse use larger bandwidths. HIFI-AKDE results in a fine resolution in dense regions and a lower resolution in sparser areas of the matrix. In order to speed up the computation of *h_i,j_*, we use a precomputed cumulative matrix, *cumRC*, where

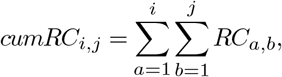

which allows the calculation of *cov_i,j_* in constant time:

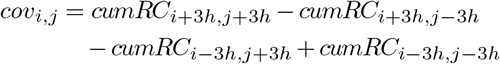

#### Markov Random Field Estimation (HIFI-MRF)

A Markov Random Field (MRF) describes a set of random variables interconnected via a lattice of dependencies. Let us denote by 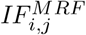 the IF value we aim to estimate at position (*i,j*). We model dependencies between neighboring cells using a log-normal distribution:

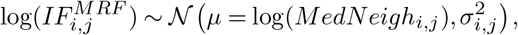

where *MedNeigh_i,j_* is the median of the eight *IF^M RF^* cells surrounding cell (*i,j*). We chose to model this dependency using the median instead of the mean of the neighbors because it allows for sharper transitions regions such as TAD boundaries. The value of 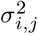 is set to *α* · log(*M edNeighb_i,j_*). *α* determines the level of dependency between cells. The value used here (*α* = 0.2) was chosen via grid search to maximize the likelihood of a left-out subset of the training data (validation set).

We model the dependency between the observed read count *RC_i,j_* and the estimated true IF value 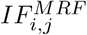 using a Poisson distribution:

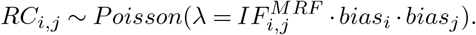

HIFI also supports the use of a negative binomial distribution to model *RC* from *IF*, to allow for increased dispersal of *RC* values via a user-defined multiplier of variance. We then seek the matrix *IF^M RF^* that maximizes Pr[*RC,IF^M RF^*] = Pr[*IF^M RF^*] · Pr[*RC* | *IF^M RF^*]. We first initialize the *IF^M RF^* matrix using the output of the HIFI-AKDE algorithm. We then optimize *IF^M RF^* using iterated conditional modes (ICM) (48) algorithm. Each iteration involves revising the value of each entry 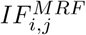 so as to maximize the joint probability of *IF^M RF^* and *RC*. Revising the value of 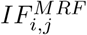 only alters the probability calculation at position *i,j* and the 8 neighboring cells (because their median may have changed), and thus probability calculations can be limited to that portion of the matrix. Because of the use of the median (rather than the mean), the joint probability function is not differentiable. Instead, the update to 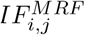 is performed by grid search over a small range of multiplicative factors. Convergence is usually achieved in 5-10 iterations over the entire matrix.

Despite using the median rather than the mean to model inter-cell dependencies, some bleed-in effect is observed at TAD boundaries. To prevent those, we designed an approach where the *nRC* matrix is first scanned to identify sharp horizontal or vertical transitions characteristic of TAD boundaries. Horizontal boundaries are defined by a row index *i* and a pair of column indices *j* and *j′*, and will be set if the distribution of *nRC* values in *nRC*_*i*,*j*…*j*′_ differs significantly from that in *nRC*_*i*+1,*j…j*′_, as determined by a Kolmogorov-Smirnov test. More precisely, boundaries are set greedily, starting with the most significant boundary matrix-wide, and iteratively adding more boundaries, provided they do not overlap previously set boundaries, until the KS statistic falls below a user-defined threshold (the value of 1.5 was used here). Vertical boundaries are symmetrical to horizontal boundaries. Boundaries are then used in the HIFI-M RF model to prevent certain neighbors from contributing to the neighborhood median of a given cell. Specifically, cells (*i′,j′*) that sit on the opposite side of a boundary from cell (*i,j*) are excluded from the neighborhood of (*i,j*).

#### Output matrices

HIFI can either produce a normalized or non-normalized output. Normalized outputs are produced by the approaches described until now. Non-normalized outputs are obtained as 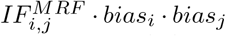. In this manuscript, normalized outputs were used throughout, except for the cross-validation experiment.

#### Conversion between fixed and restriction fragment resolutions

HIFI operates at RF resolution, whereas other approaches operate at fixed resolutions (e.g., 5, 25, or 50 kb). To convert from fixed to RF resolution, we tested three alternatives: the ‘direct’ scheme, which assigns each restriction fragment the frequency of the fixed bin that their 3’ end resides within; (ii) the ‘counts’ scheme, which divides a fixed bin’s interaction frequency by the number of 3’ fragment ends found within it and then assigns this value to each of those fragments; (iii) the ‘weighted’ scheme, which determines the proportion of each fragment that overlaps with a given fixed bin, then assigns each fragment its relative proportion of that bin’s interaction frequency. The ‘direct’ conversion scheme was found to be the most robust and perform more consistently across datasets, and is the one that was used throughout the paper.

### Alternative approaches

The source code for HiC-Plus (17) was obtained from https://github.com/zhangyan32/HiCPlus. Models were trained on Hi-C data from chromosomes 1-8 at 10 kb resolution, within a range of 2 Mb, as recommended. Input and target contact frequencies were obtained from input set and test *RC* matrices, respectively. Models were provided 100 epochs (10 times more than recommended) to converge while ensuring overfitting did not occur.

HMRFBayes (15) was obtained from http://www.unc.edu/~xuzheng/HMRFHiCFast/tutorial.php.

The HM RFBayes program was provided the observed and expected contact frequencies for paired restriction fragments within 1 Mb bins along chromosomes 9-X, where the expected contact frequency was calculated as follows:

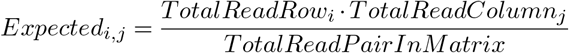

### Directionality index and TAD boundary prediction

The directionality index (DI) was first described by Dixon et al. (2012) (6) to detect directionality bias for interactions across a Hi-C IF matrix. For RF *i*, the DI is usually calculated as follows:

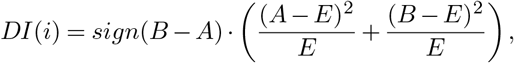

where *A* = ∑_*i−δ≤j<i*_ *IF_j,i_, B* = ∑_*i<j≤i+δ*_ *IF_i,j_*, 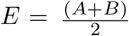, and *δ* is set to 500 kb. Due to the low coverage at RF-resolution Hi-C data, the DI formula yields very noisy predictions. We thus used the following modified version:

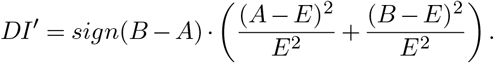

This modification transforms terms present in the right parentheses to relative rates and helps to scale the magnitude of the DI. TAD boundaries are defined as RFs whose *DI′* value is a local maximum or minimum in a window of 21 RFs (51 for MboI analyses) centered around it. In the case of the fixed binning (b=16) analysis, only RFs at the center of their bin are considered. Due to their low coverage, regions within 2 Mb of a centromere or telomere were excluded. TAD boundaries are then sorted by their absolute DI values and the top 5,000 and 25,000 boundaries were selected for HindIII and MboI RF-resolutions, respectively.

### Data sources and pre-processing

The following Hi-C data sets were used. From Rao et al. (2014) (23), GM12878 with HindIII and MboI digest (GEO:GSE63525); From Dixon et al. (2012) (6), mESC with HindIII digest (GEO:GSE35156). For 5C comparisons, the following data sets were used: From Smith et al. (2016) (28): GM12878 with HindIII digest (GEO:GSE75634); From Nora et al. (2012) (7): mESC with HindIII digest (GEO: GSE35721). For comparisons to ChIA-PET, the following data sets were used: From Tang et al. (2015) (31), CTCF-mediated contacts (GEO:GSM1872886) and RNAPII-mediated contacts (GEO:GSM1872887). From Fullwood et al. (2009) (30): RAD21-mediated contacts (GEO: GSM1436265; replicates averaged). Paired-end tag clusters were binned to hg19 HindIII RFs to ensure comparability with other datasets. Enhancer-promoter (EP) pairs from Thurman et al. (2012) (33) were obtained from: ftp://ftp.ebi.ac.uk/pub/databases/ensembl/encode/integration_data_jan2011/byDataType/openchrom/jan2011/dhs_gene_connectivity/genomewideCorrs_above0.7_promoterPlusMinus500kb_withGeneNames_32celltypeCategories.bed8.gz. Enhancers and promoters were then binned to their respective RFs.

ChIP-seq data from ENCODE (34) and ChromHMM (43) predictions were downloaded from the UCSC Genome browser (49) and binned to HindIII and MboI RFs. For ChromHMM (Fig. 5C and 5G), only states 1 and 4 were used (to reduce redundancy). CTCF motifs and orientation were identified in a similar manner to Fundenberg et al. (2016) (50) using HOMER (51) and the ‘CTCF_known1’ PWM (52). CTCF peaks were then assigned assigned directionality based on overlapping CTCF motifs. CTCF peaks, identified by ChIP-seq, were assigned forward or reverse strand orientations based on overlapping CTCF motifs. If both orientations were found to reside within a peak, then one orientation would be randomly chosen. Peaks with no orientation or overlapping CTCF motif were discarded.

## ACKNOWLEDGEMENTS

This work was funded by a NSERC Discovery grant to M.B. and a CIHR grant (MOP-142451) to J.D. Authors would like to thank X.Q. David Wang, Maia Kaplan, Rola Dali, Jack Guo, Yanlin Zhang, Jérôme Waldispühl, Derek Ruths, Michael Hallett, Alessandro Bonetti, Michael Hoffman, Michiel de Hoon, and Nicole Francis for useful discussions during the development of this project and manuscript.

---

**Supplementary Figure S1.**
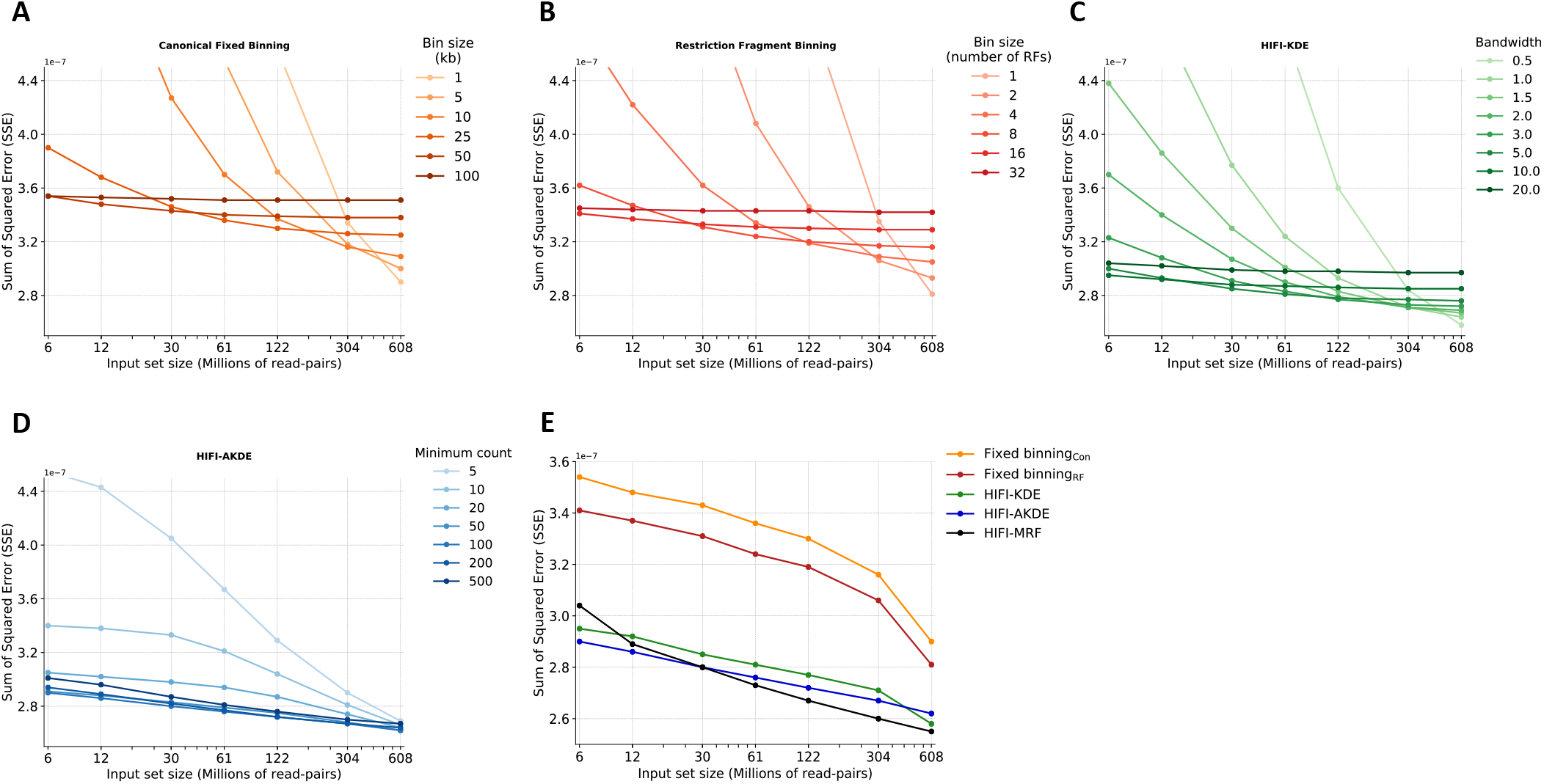
Cross-validation of fixed-binning and HIFI methodologies. **A**) [Reproduced from Fig. 1B, to facilitate comparison.] Cross-validation error for canonical fixed-binning approaches, for different bin sizes, as a function of coverage. **B**) Same analysis, performed on fixed-binning algorithm where bin size is expressed in terms of the number of restriction fragments rather than in kb. **C**) Cross-validation error for HIFI-KDE, at different bandwidth values. **D**) Cross-validation error for HIFI-AKDE, at different Minimum Count values. **E**) Comparison of most accurate parameter sets at various input set sizes for all inference methodologies (based on SSE). We observe that HIFI-MRF outperforms all other approaches described for most input set sizes.

**Supplementary Figure S2.**
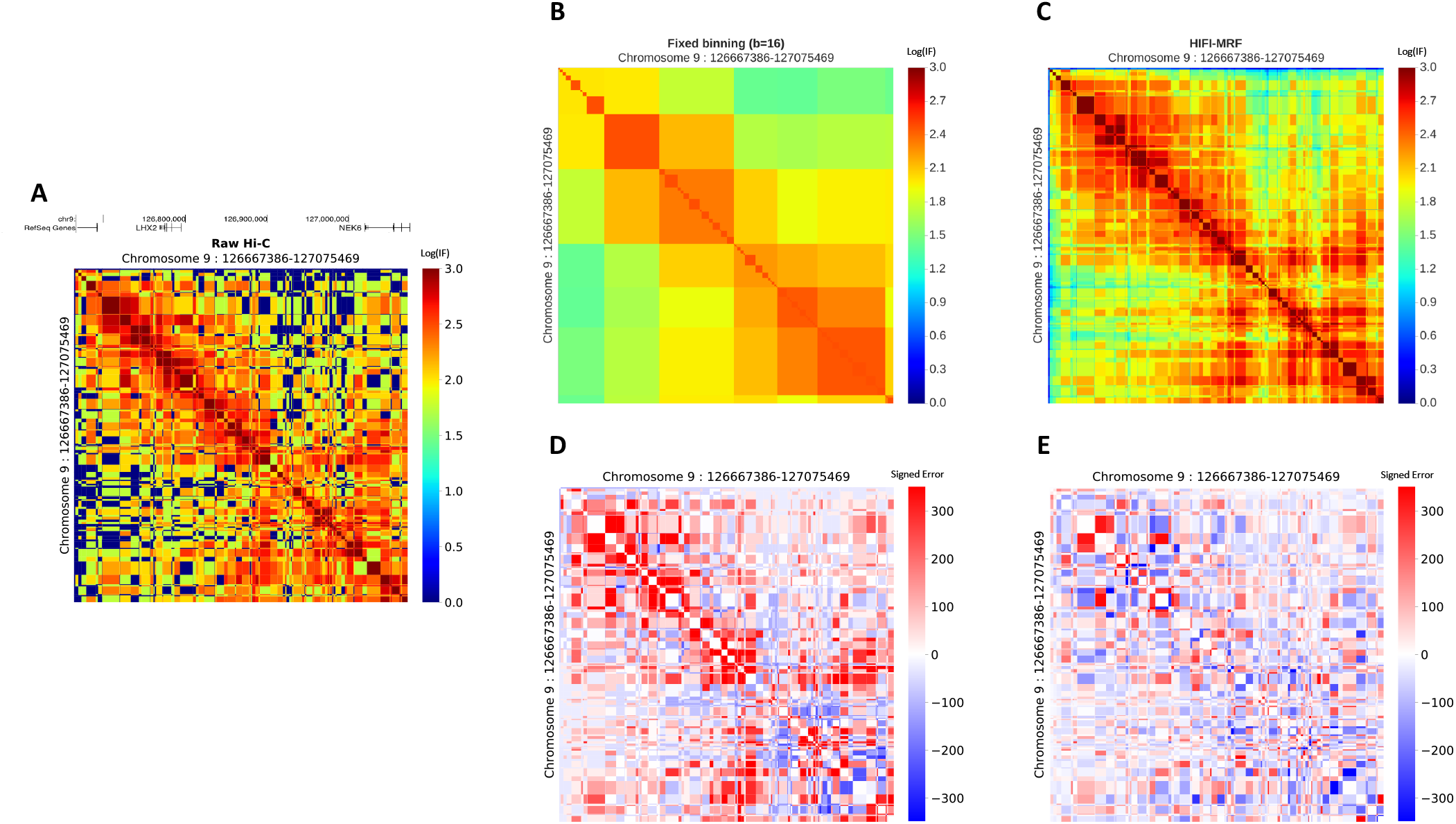
Fixed-binning approach vs. HIFI-MRF. **A**) Raw Hi-C IF matrix for the NEK6 locus at HindIII RF-resolution. Inferred Hi-C IF matrices resulting from fixed-binning (b=16 RFs) and HIFI-MRF approaches is shown in (**B**) and (**C**), respectively. Signed error matrices (resulting from the subtraction of the raw Hi-C matrix [**A**] by either [**B**] or [**C**]) are shown for fixed binning (**D**) and HIFI-MRF (**E**). A noticeable reduction in error is observed for the HIFI-MRF signed error (**E**) when compared to the fixed-binning (**D**).

**Supplementary Figure S3.**
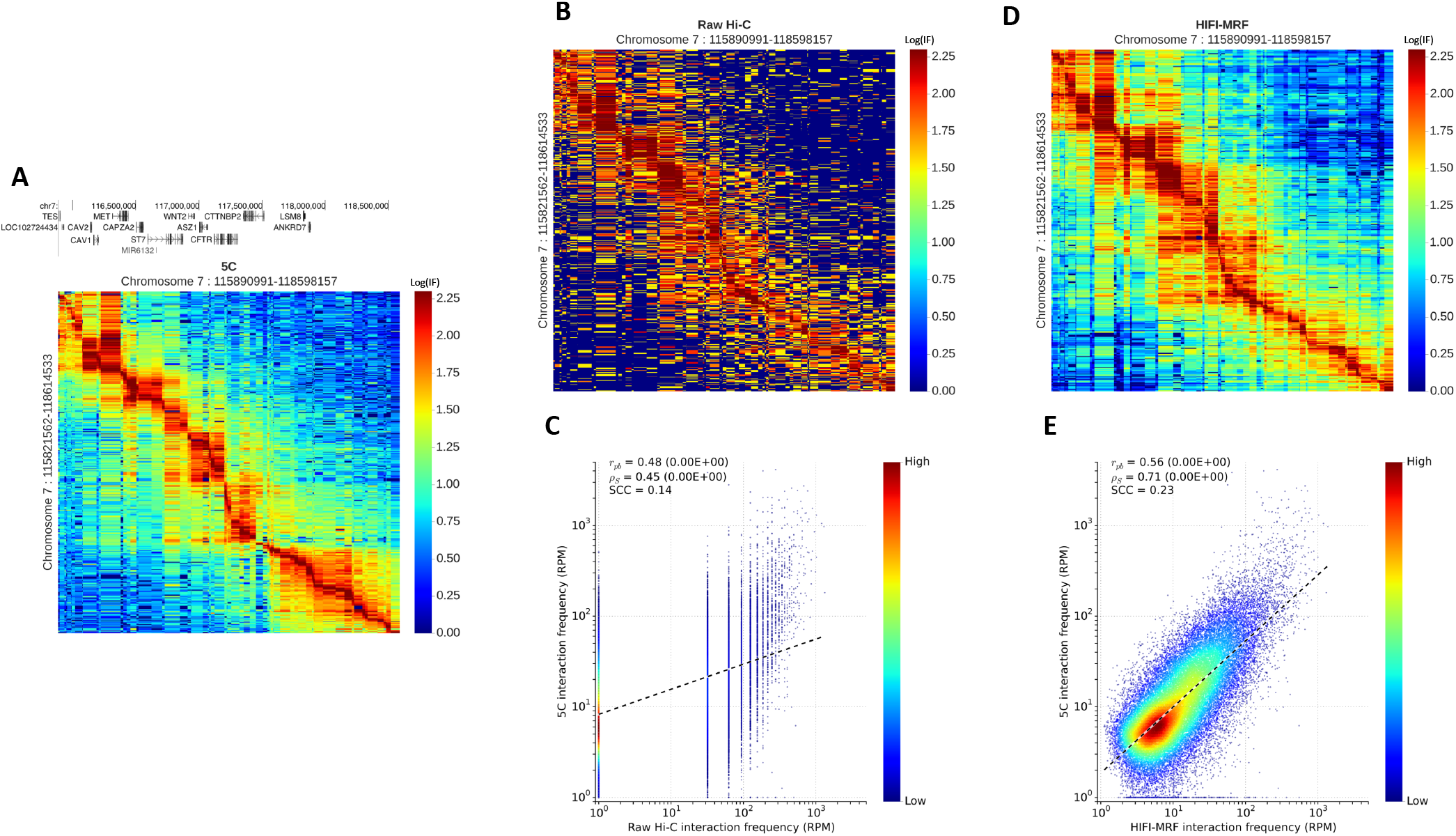
Recapitulation of 5C observations by HIFI-MRF (GM12878-chr7 example). 5C data for chr7:115890991-118598157 in GM12878 cells (**A** (28)). (**B**) Raw HiC data (23) for the corresponding region. (**C**) Correlation of raw HiC data with 5C data for this region (Pearson *r_pb_* = 0.48, p-value< 10^−16^; Spearman *ρ_s_* = 0.45, p-value< 10^−16^); SCC = 0.14). (**D**) HIFI-MRF inferred IFs for the same region. (**E**) Correlation of HIFI-MRF inferred IFs with 5C data for this region (Pearson *r_pb_* = 0.56, p-value< 10^−16^; Spearman *ρ_s_ =* 0.71, p-value< 10^−16^); SCC = 0.23). Also observe how HIFI-MRF processed data displays TADs and a decay constant profile similar to those observed by 5C.

**Supplementary Figure S4.**
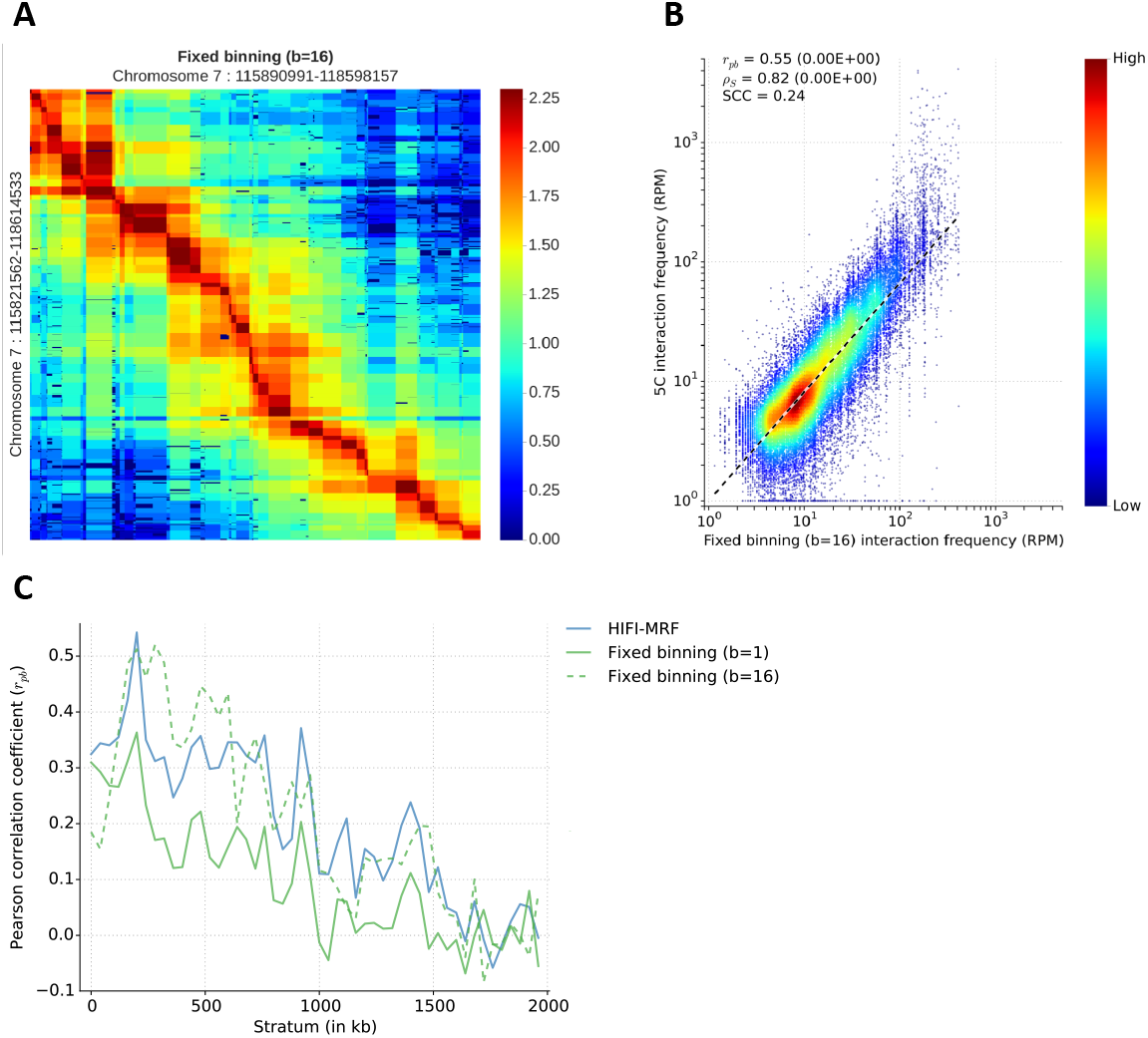
Fixed-binning recapitulation of 5C contacts (GM12878-chr7 example). (**A**) Fixed binning (b = 16) analysis of Hi-C data for GM12878-chr7 RF-pairs (28). (**B**) Comparison of observed 5C and fixed-binning (b = 16) Hi-C values for the region (Pearson *r_pb_* = 0.55, p-value< 10^−16^; Spearman *ρ_s_* = 0.82, p-value< 10^−16^); SCC = 0.24). (**C**) Distribution of Spearman *ρ_s_* values by genomic distance for Hi-C contacts resulting from HIFI-MRF and fixed binning. Both HIFI-MRF and fixed-binning perform similarly for this example.

**Supplementary Figure S5.**
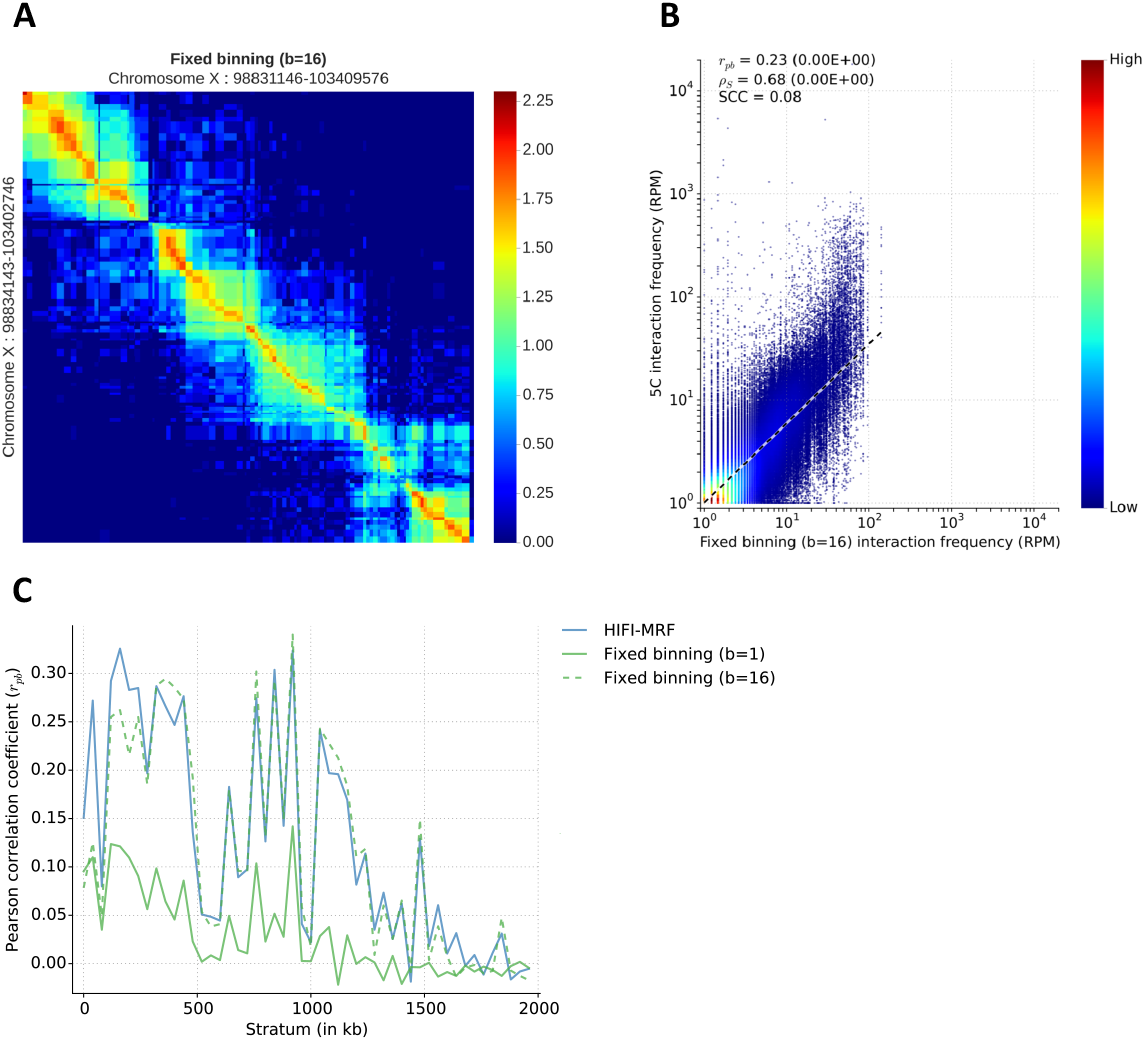
Fixed-binning recapitulation of 5C contacts (mESC Xist locus). **A**) Fixed binning (b = 16) analysis of Hi-C data for a 4.5 Mb region surrounding Xist in mESC. **B**) Comparison of observed 5C (7) and fixed-binning (b = 16) Hi-C values for the region (Pearson *r_pb_* = 0.23, p-value< 10^−16^; Spearman *ρ_s_* = 0.68, p-value< 10^−16^); SCC = 0.08). **C**) Distribution of Spearman *ρ_s_* values by genomic distance for Hi-C contacts resulting from HIFI-MRF and fixed binning. Overall, due to the low SCC (29) value and similarity in distributions of panel **C**, most of the improvement by either method is driven by distance dependencies between contacts.

**Supplementary Figure S6.**
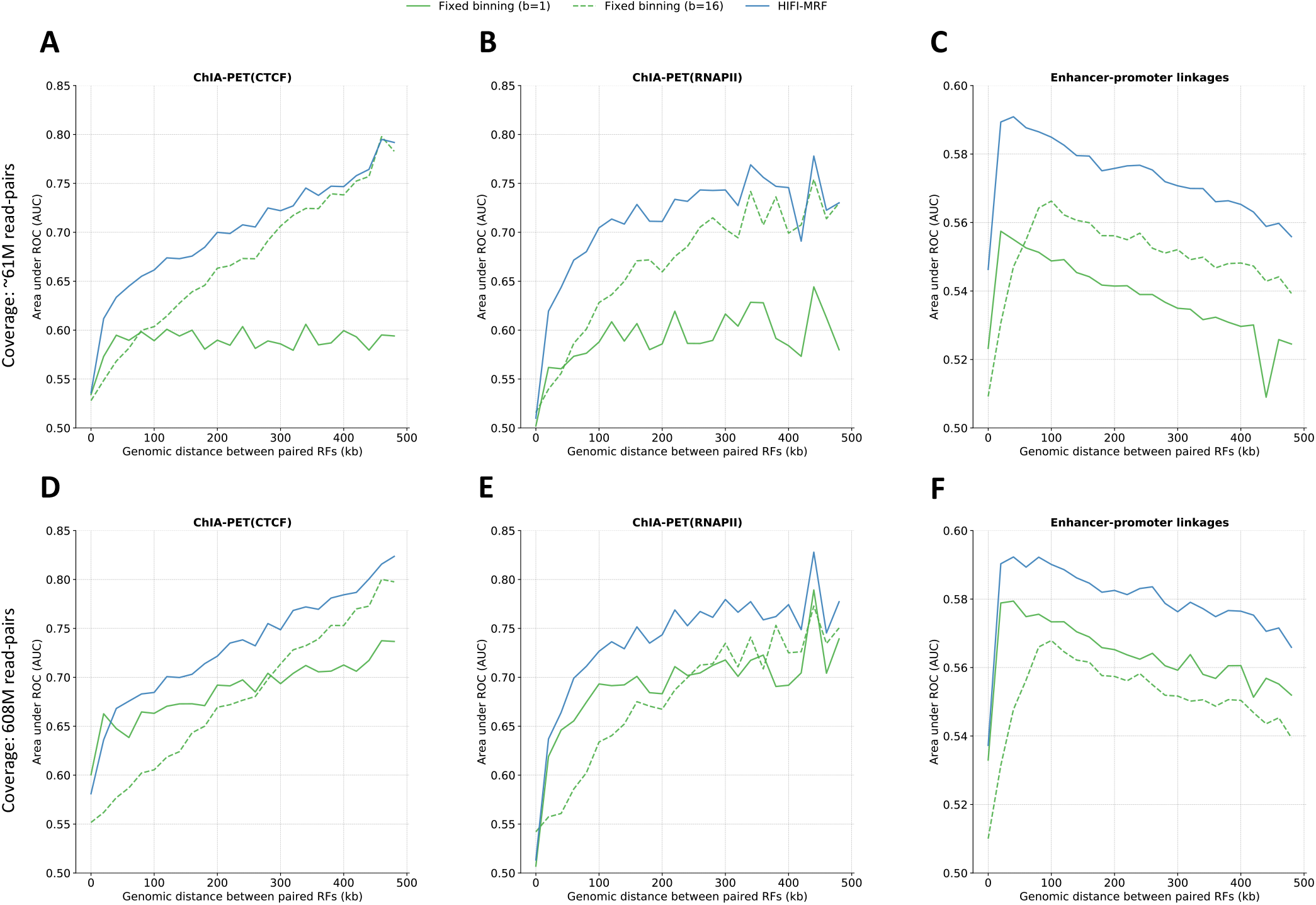
Positive/negative RF contact delineation analysis (genome-wide). Repetition of the analysis shown in Fig. 4, but this time for the whole genome. See caption of Fig. 4

**Supplementary Figure S7.**
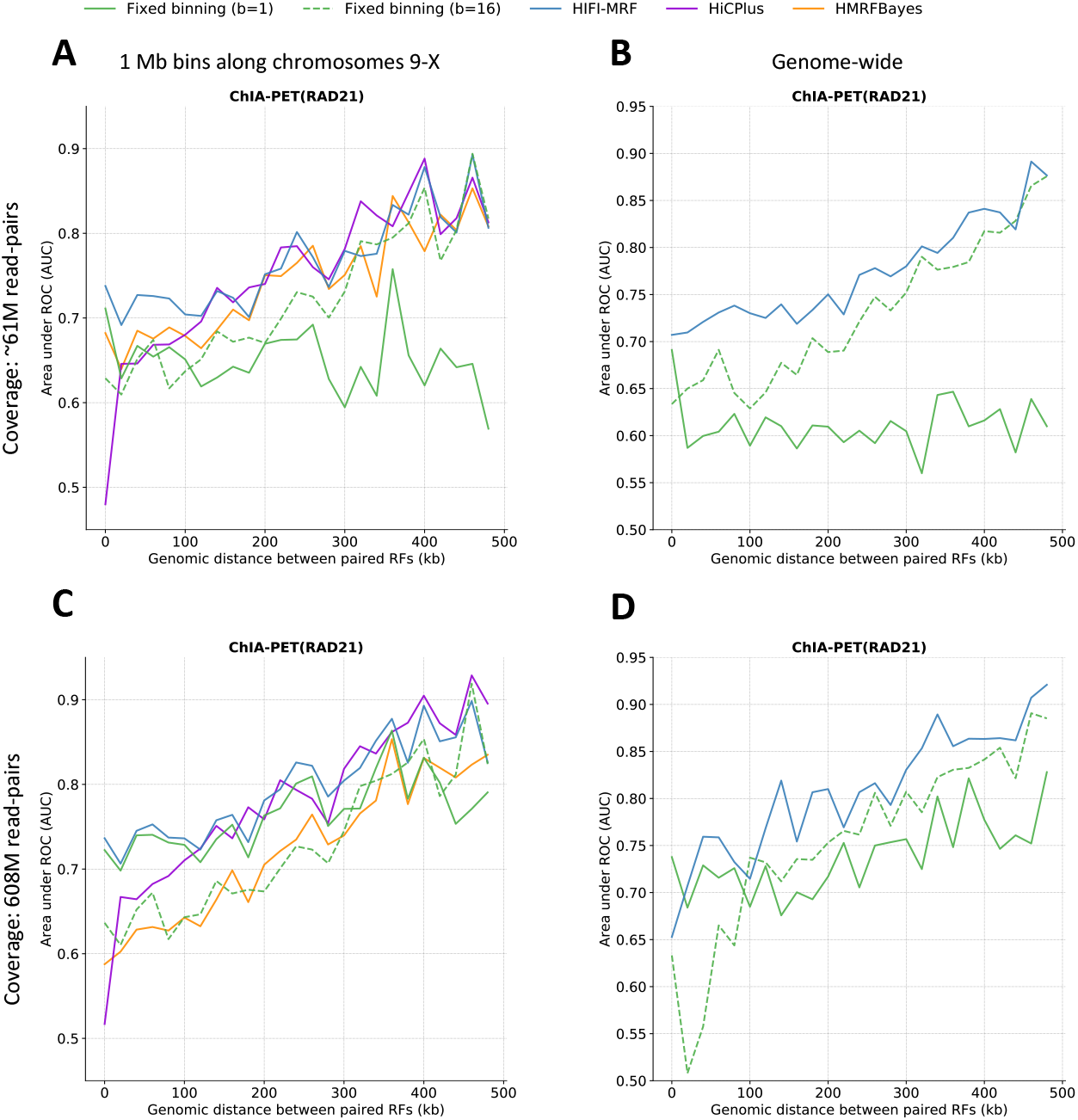
Positive/negative RF contact delineation analysis (RAD21). Repetition of the analysis presented in Fig. 4, this time for ChIA-PET RAD21 data (32). HIFI-MRF is found to provide more accurate (based on AUROC) predictions of RF-pair classification (positive vs. negative) compared to other inference methods for contacts located less than 100 kb apart, but is on par with HiCPlus and HMRFBayes at larger distances.

**Supplementary Figure S8.**
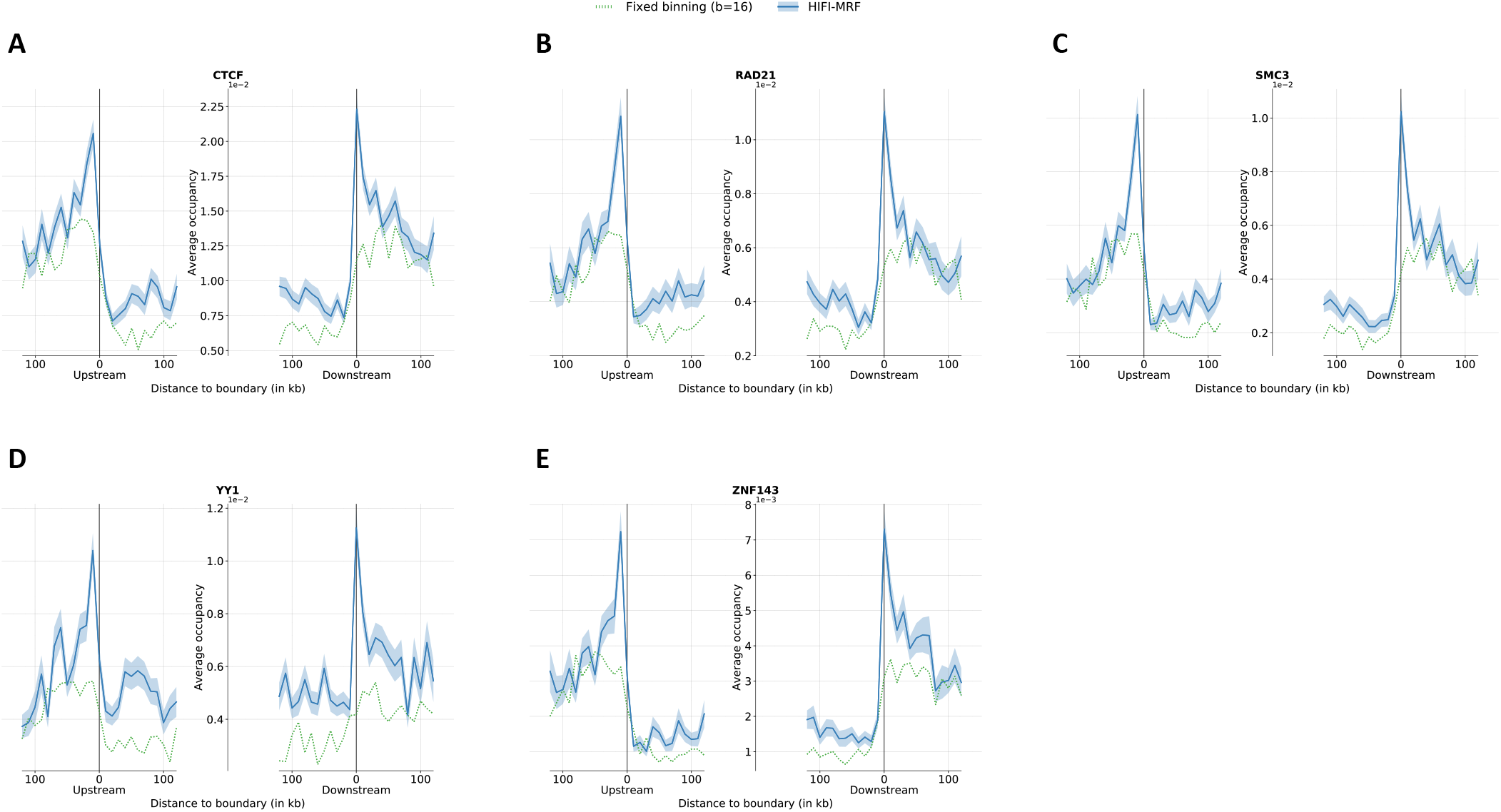
GM12878-HindIII RF-resolution TAD boundary occupancy by architectural proteins. Architectural proteins CTCF (**A**), RAD21 (**B**), SMC3 (**C**), YY1 (**D**), and ZNF143 (**E**) are strongly enriched near TAD boundaries, followed by a significant depletion in occupancy within TADs.

**Supplementary Figure S9.**
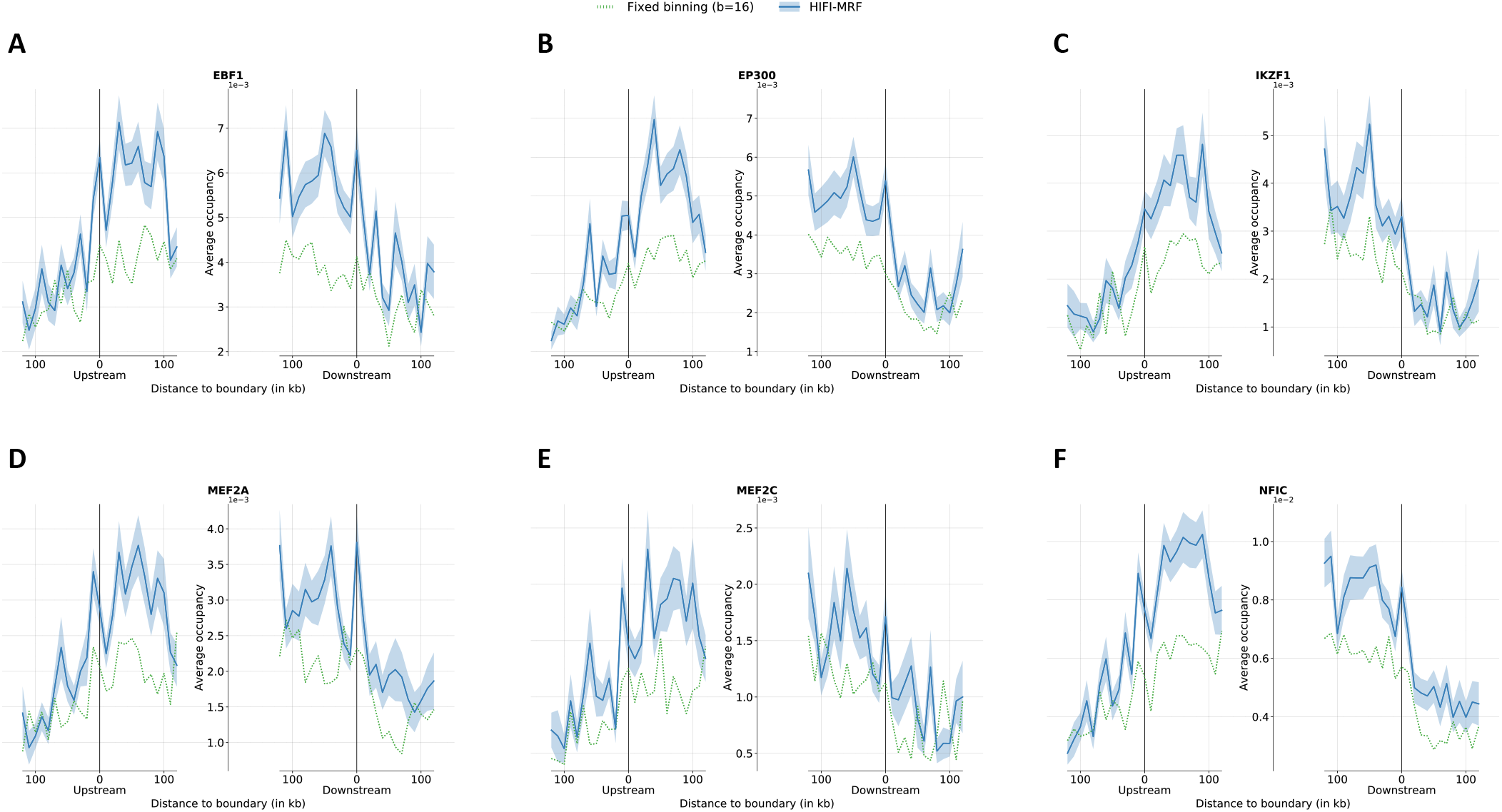
GM12878-HindIII RF-resolution within-TAD enrichment by selected transcription factors. Transcription factors EBF1 (**A**), EP300 (**B**), IKZF1 (**C**), MEF2A (**D**), MEF2C (**E**), and NFIC (**F**) show an enrichment within TADs, compared to outside of TAD boundaries.

**Supplementary Figure S10.**
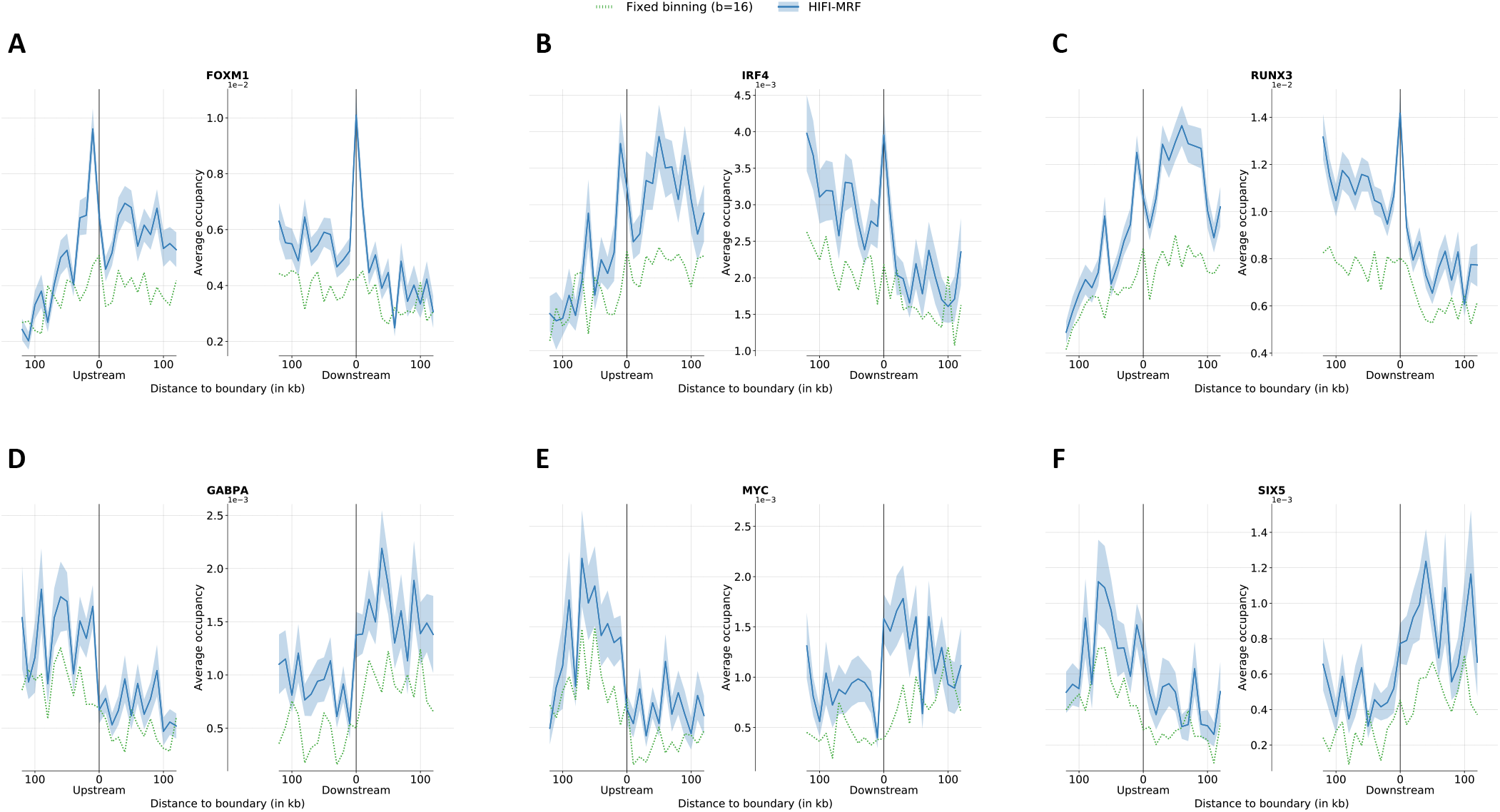
GM12878-HindIII RF-resolution TAD boundary occupancy of selected transcription factors. Transcription factors FOXM1 (**A**), IRF4 (**B**), and RUNX3 (**C**) exhibit a strong enrichment specific to TAD boundaries, in addition to an enrichment within TADs. On the opposite, transcription factors GABPA (**D**), MYC (**E**), and SIX5 (**F**) show a minor depletion within TADs compared to outside of TADs.

